# Effects of arginine on the interfacial behavior of proteins

**DOI:** 10.1101/2025.09.07.674647

**Authors:** Peyman Dastyar, Harsa Mitra, Caitlin V. Wood, Yu-Jiun Lin, Yu Tian, Arezoo M. Ardekani

**Affiliations:** School of Mechanical Engineering, Purdue University, 585 Purdue Mall, West Lafayette, IN 47907, United States of America; Merck & Co., Inc., Rahway, NJ 07065, United States of America

**Keywords:** Interfacial rheology, Arginine, Fusion protein, Aggregation, Protein-protein interactions, Amino acid, Protein adsorption

## Abstract

**Hypothesis:** Protein adsorption at fluid–fluid interfaces can induce conformational rearrangements and promote aggregation. Arginine, as an amino acid is known to attenuate protein–protein interactions and lower the viscosity of protein formulations. Given its surface-active properties, we hypothesize that arginine may be effective in limiting aggregation at fluid–fluid interfaces.

**Experiments:** Interfacial rheology of Etanercept (Enbrel), Immunoglobulin G1 (IgG1), and Bovine Serum Albumin (BSA) was characterized across a range of concentrations, with and without L-arginine Hydrochloride. The interfacial response was examined in both linear and nonlinear regimes. *ζ*-potential was also measured for selected formulations to evaluate the relationship between electrostatic characteristics and the rheological outcomes.

**Findings:** Based on both linear and nonlinear interfacial rheology results, arginine generally weakened protein–protein interactions; however, the effect depends on protein identity and concentration. In arginine-containing formulations, Enbrel and BSA displayed only minor variation in interfacial viscoelasticity between 0.01 and 1 mg/mL, particularly during prolonged aging; in contrast, IgG1 showed larger concentration-dependent differences. These results strengthen predictive frameworks for interfacial stability and guide the rational design of protein–amino acid biopharmaceutical formulations.

**Graphical Abstract:** 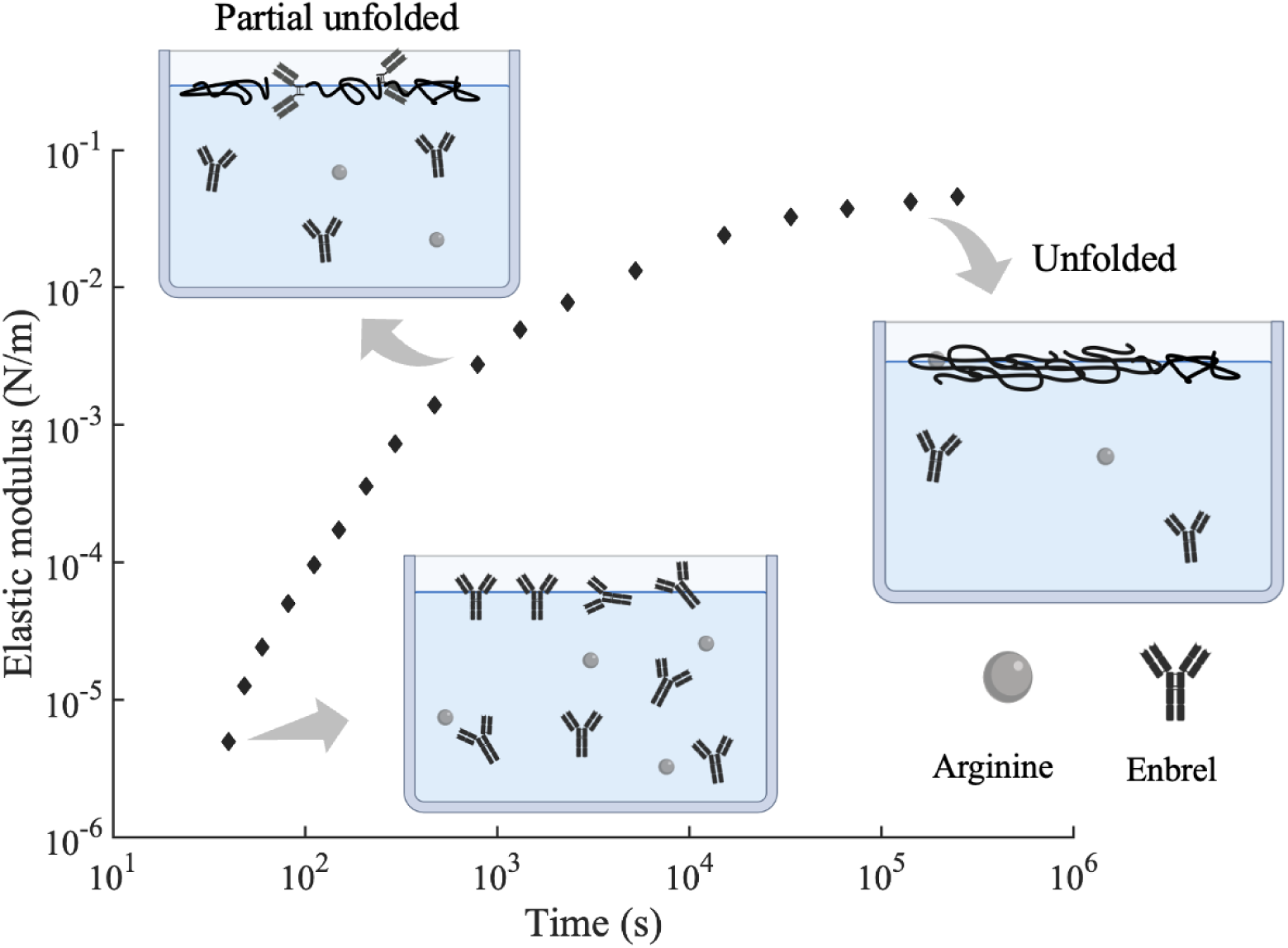

## 1. Introduction

Over the past few decades, monoclonal antibodies (mAbs) have been extensively utilized in the pharmaceutical industry to treat a variety of diseases, including cancer, respiratory conditions, and autoimmune disorders [1, 2]. These biologic therapeutics are distinguished by their precise targeting of specific cells or proteins and minimal side effects, which place them among the fastest-growing pharmaceutical products [3]. In this study, we focus on Enbrel, or Etanercept, a type of fusion protein with a similar structure to mAbs but with key differences that affect its behavior. Fusion proteins are created by joining two protein domains, giving them multifunctional properties derived from the parent molecules [4]. For instance, Enbrel consists of two molecules of the 75 kDa fragment of the human tumor necrosis factor *α* (TNF*α*) receptor fused to the Fragment crystallizable (Fc) portion of immunoglobulin IgG1 [5, 6, 7, 8]. The Fc fragment extends the half-life of the fusion protein within the vascular system, resulting in a more prolonged biological impact [6]. On the other side, the TNF*α* receptor binds to TNF*α*, making it effective in treating various inflammatory diseases such as rheumatoid arthritis (RA), juvenile rheumatoid arthritis (JRA), psoriatic arthritis, ankylosing spondylitis, and psoriasis [5, 9].

One of the primary challenges in the manufacturing and storage of therapeutic proteins is their inherent instability in solutions, which can lead to undesirable immune responses and diminished therapeutic efficacy [10, 11]. Protein instability may manifest as conformational changes from native to non-native states or self-association of protein molecules, influenced by various factors such as solution pH, protein interactions, concentration, and temperature. To address these issues, the pharmaceutical industry often employs excipients—additives such as salts, sugars, amino acids, and surfactants—in therapeutic formulations. Arginine, an abundant amino acid in protein structures [12, 13, 14], is widely utilized to facilitate the refolding of recombinant proteins, reduce non-specific protein interactions with solid surfaces, and mitigate protein-protein interactions [15, 16, 17]. Due to its unique ability to suppress protein aggregation without causing denaturation [18, 19], arginine is an integral component in medical formulations that primarily contain Enbrel. However, the complex structure of proteins and the multifaceted effects of arginine on protein-protein interactions pose significant challenges in discerning and predicting the precise mechanisms by which arginine influences protein behavior.

Arginine is capable of interacting with protein molecules in multiple ways due to the presence of four functional groups in its structure: i) a guanidinium group, ii) an amino group, iii) a carboxylate group, and iv) a methylene group [20]. The guanidinium group of arginine is positively charged [21], allowing it to form cation-*π* interactions with aromatic residues of proteins [13, 22, 23, 12]. Additionally, this group exhibits a strong propensity for hydrogen bond donation [12], which enhances arginine’s binding affinity to protein surfaces. The presence of the guanidinium group is crucial for binding to protein surfaces and mitigating electrostatic and hydrophobic interactions among protein molecules. Furthermore, the carboxylate residue within arginine is known to inhibit interactions involving its guanidinium group and proteins [22], while the methylene group contributes to arginine’s hydrophobic properties [12]. Arginine’s role in protein aggregation is further influenced by the electrostatic screening effects of its salts [24]. As a positively charged molecule, arginine binds to anionic species in the commercially available chemical formulations, functioning similar to a salt in protein solutions. This interaction can modulate electrostatic forces, either diminishing or enhancing attractive or repulsive interactions among proteins. The versatility of arginine’s chemical structure allows it to participate in diverse mechanisms of interaction with proteins, thereby potentially influencing protein-protein interactions based on the specific chemical makeup of the proteins involved and the nature of the arginine salt’s anionic component [11].

The extensive body of literature on protein-protein interactions and their implications for the mechanistic properties of solutions has concentrated significantly on understanding the factors influencing protein solution viscosity. Previous studies reveal that protein interactions can arise from various factors, leading to the aggregation of protein molecules and consequently affecting the bulk viscosity of the solution. Arginine, as an additive, has been identified as an effective agent in reducing viscosity; however, its impact on protein stability varies under different conditions. Dear et al. [25] examined the effects of various excipients on the viscosity of a mAb, and the findings showed the reducing effects of ArgHCl in viscosity. Their analysis suggested that arginine mitigates hydrophobic interactions and electrostatic attractive forces by increasing positive charges on the protein surface, thereby reducing local interactions between cations and anions. Similar results were reported by Hu et al.[22], who demonstrated that arginine was also successful in preserving the monomer content in the solution. Additionally, other investigations [22] proposed that electrostatic forces are the dominant factor in protein-protein interactions; however, arginine also influences other interactions such as van der Waals forces, hydrogen bonding, *π*-*π* stacking, and hydrophobic interactions. Further research by Chowdhury et al.[26], Inoue et al.[27], Borwankar et al.[21], Wang et al.[24], and Kim et al. [28] also highlights the role of arginine in attenuating protein interactions and reducing the viscosity of various monoclonal antibodies.

Despite numerous studies demonstrating the anti-aggregation properties of arginine in protein systems, conflicting findings have also been reported. For instance, Borzova et al. [29] investigated the time-dependent aggregation of heat-induced BSA in the presence of arginine and found that it increased the hydrodynamic diameter of the aggregates in solution. They further noted that the effect of arginine on aggregate-aggregate assembly depends on its concentration. In another study focusing on human immunoglobulin G (IgG) aggregation [18], arginine was found to exhibit dual effects: while it facilitated protein unfolding, it also suppressed aggregation, indicating its complex impact influenced by various factors.

Based on the aforementioned survey, while numerous studies have examined arginine’s impact on the rheological properties of proteins, most have focused on its role in the bulk solution, particularly in relation to viscosity reduction [21, 24, 26, 27]. Systematic investigations of how arginine (or, more generally, amino acids) modulates protein self-association at fluid–fluid interfaces, where behavior can differ fundamentally from the bulk, are scarce. Prior work has characterized interfacial behavior of protein systems [30, 31] and the inhibition of interfacial self-association by surfactant-active additives [1, 32, 33]; however, the specific interplay between proteins and amino acids at fluid–fluid interfaces has rarely been addressed. Moreover, molecular-level insights into protein–protein interactions at such interfaces, especially for fusion proteins whose structures differ significantly from typical monoclonal antibodies, are largely absent from the literature. In this study, we investigate the interfacial rheology of Enbrel, IgG1, and BSA in the presence of arginine to elucidate the mechanisms of interaction between arginine and protein molecules and how it influences protein adsorption and network formation at the air-water interface. Comparing these three types of proteins provides deeper insight into the chemical phenomena underpinning molecular interactions.

## 2. Materials and Methods

### 2.1. Sample preparation

Enbrel (Etanercept), utilized in this study, was supplied by Immunex Corporation and provided in pre-filled syringes. Each syringe contains 0.98 mL of a solution consisting of 50 mg/mL Enbrel, 1% sucrose, 100 mM Sodium Chloride, 25 mM L-arginine Hydrochloride, and 25 mM Sodium phosphate, with a pH of 6.3 *±* 0.2. Enbrel comprises 934 amino acids and has a molecular weight of approximately 150 kDa. Human IgG1 Kappa, purified from human myeloma serum, was procured from Southern Biotech. This protein has a molecular weight of 146 kDa and was supplied at a concentration of 0.5 mg/mL in a 1 mL volume, used without further purification. Bovine Serum Albumin (BSA) (purity *≥* 98%, MW 66 kDa) and L-Arginine Hydrochloride were prepared from Sigma Aldrich. Additionally, all solutions were prepared in 25 mM phosphate saline buffer solutions with a pH of 6.5. Sodium phosphate monobasic monohydrate and Sodium phosphate dibasic heptahydrate, also purchased from Sigma Aldrich, were used to prepare the buffer solution at the specified pH. We examined four concentrations of 0.001, 0.01, 0.1, and 1 mg/mL for Enbrel and BSA, and three concentrations of 0.001, 0.01, and 0.1 mg/mL for IgG1. Furthermore, the behavior of these proteins was investigated in both phosphate buffer solutions without arginine and in the presence of 25 mM L-arginine Hydrochloride. Although therapeutic formulations of monoclonal antibodies and fusion proteins are typically designed at high protein concentrations, prior investigations into protein adsorption at fluid–fluid interfaces have consistently shown that interfacial saturation occurs at much lower concentrations [34]. Specifically, saturation concentrations for monoclonal antibodies are reported in the range of 0.1–1 mg/mL [35], while for BSA, saturation is observed at even lower concentrations, typically between 0.001–0.01 mg/mL [36]. Therefore, we considered the selected concentration range in this study to be sufficient to capture the full interfacial adsorption behavior and viscoelastic transition from dilute to saturated states.

We conducted dialysis on the Enbrel solutions to eliminate any excess components. The procedure involved injecting the solution into a dialysis cassette obtained from ThermoFisher Scientific (Slide-A-Lyzer™ G3 Dialysis Cassettes with a 10 kDa molecular weight cut-off and a 3 mL sample volume). The cassette was then immersed in 1 L of 25 mM phosphate buffer (prepared with Sodium phosphate monobasic monohydrate and Sodium phosphate dibasic heptahydrate, and deionized distilled water) at pH 6.5. The dialysis process was carried out by exchanging the buffer after the first 8 hours. The second exchange occurred 8 hours later. Subsequently, the buffer was changed twice more, each time after 24 hours. Dilute solutions of the withdrawn sample were subsequently prepared, and the concentration of the dialyzed protein was determined by measuring the average concentration of these dilute solutions using a spectrophotometer (Thermo Scientific NanoDrop 2000C).

### 2.2. Interfacial Rheology - linear viscoelastic regime (LVE)

The interfacial rheology tests were conducted using the Discovery HR-2 from TA Instruments. The selected geometry was the Double Wall Ring (DWR), which consists of a thin ring positioned at the fluid interface. This geometry is designed to have a high contact perimeter-to-area ratio [37], ensuring that the bulk contribution to the measured stress values is negligible. Each test required 17-18 mL of liquid, and the temperature for all tests was consistently maintained at 6°C. Rheological tests for selected formulations—including BSA 0.1 mg/mL, BSA 1 mg/mL, Enbrel 0.001 mg/mL, Enbrel 0.1 mg/mL with arginine, and IgG1 0.01 mg/mL with arginine—were repeated three to five times. The average coefficient of variation in the measured complex modulus at 30,000 seconds across these replicates was 11.2 %. Viscoelastic measurements were performed at a 2% strain amplitude and 1 rad/s frequency, falling within the linear viscoelastic regime (LVE) for all sample interfaces. The sequence of tests began with positioning the geometry at the fluid interface, followed by a time-sweep test conducted for 10 hours. Subsequently, frequency-sweep (at a strain amplitude of 2%) and amplitude-sweep tests were performed. The relaxation and flow-sweep tests were conducted separately after the 10-hour time-sweep test to ensure that tests inducing non-linear deformations did not interfere with the other measurements.

### 2.3. Interfacial Rheology - Large Amplitude Oscillatory Shear (LAOS)

The strain-sweep tests were performed at a constant frequency of 1 rad/s, with the strain amplitude varying from 0.1% to 100%. During each cycle, a sinusoidal strain, represented as *γ* = *γ*_0_ sin *ωt* was applied to the interface, where *γ*_0_ and *ω* denote the strain amplitude and frequency, respectively. The resulting stress (*σ*(t)) was recorded over time. The collected data was subsequently analyzed using the MITLOAS software (for comprehensive theoretical derivations, refer to Ewoldt et al. [38]), yielding the stress as a function of deformation, which can be depicted in the form of Lissajous–Bowditch curves. The morphology of these curves elucidates the nonlinear rheological properties of the interface. In addition, in order to quantitatively assess the nonlinearity and enhance the interpretation of material behavior under varying conditions, six new parameters have been introduced [38, 39, 40], which depend on the geometry of the curves.

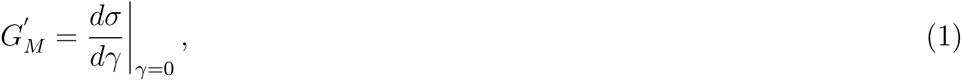

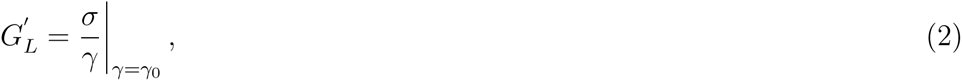

where *G*^′^ and *G*^′^ denote the minimum-strain and large-strain moduli, respectively. Likewise, the minimum-rate dynamic viscosity (*η*^′^) and largerate dynamic viscosity (*η*^′^) are defined, as shown in the following equations:

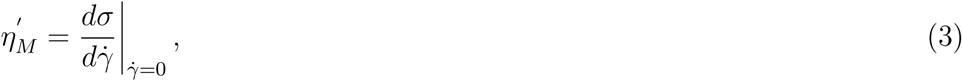

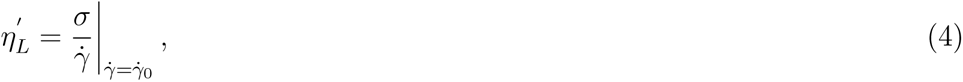

Consequently, the strain-stiffening ratio (S) and shear-thickening ratio (T) are explicitly defined by Equations 5 and 6. The values of S and T serve as indicators of the intra-cycle nonlinearity of the protein solutions’ interface. A positive S indicates strain-stiffening, suggesting increased resistance to deformation under strain, while a negative S signifies strain-softening. Similarly, a positive T value denotes shear-thickening behavior, where viscosity increases with shear rate, while a negative T value indicates shear-thinning, where viscosity decreases with shear rate.

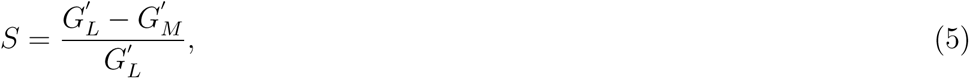

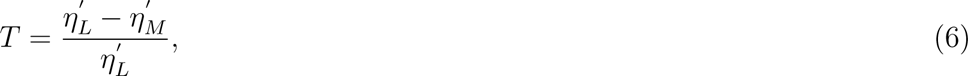

### 2.4. ζ-potential measurements

*ζ*-potential is a key parameter that quantifies the magnitude of electrostatic repulsive forces between colloidal particles, as explained by the electrical double layer theory [41]. It can be helpful in understanding the stability of colloidal particles, as higher *ζ*-potential values typically indicate greater repulsion. In this study, the *ζ*-potential was measured using a Malvern Nano ZS90 system equipped with a 4 mW, 532.8 nm red laser and DTS1070 disposable cuvettes. All measurements were performed at a precisely controlled temperature of 6 ^◦^C to ensure consistency and accuracy. Each test was repeated a minimum of three times to minimize variability, and the average values of these measurements are presented in this work. It is important to note that the samples used for these *ζ*-potential measurements were prepared independently from those used in the interfacial rheology experiments.

## 3. Results

### 3.1. Interfacial Rheology measurements

To better comprehend the behavior of the proteins and determine the linear viscoelastic regime, the viscoelasticity of the protein solutions at different concentrations, in the presence and absence of arginine, was measured in terms of strain amplitude. The frequency was kept constant at 1 rad/s across all tests, and the data for a 0.1 mg/mL concentration is shown in Figure 1. As observed, Elasticity remains almost constant at low strain amplitudes and starts to decrease at higher values. The range of strain over which the elastic modulus is constant is referred to as the linear viscoelastic (LVE) regime. In this regime, the network structure of the protein molecules is perturbed but remains intact. At higher strains, the molecular networks begin to break down and the protein entities separate from each other.

**Figure 1:**
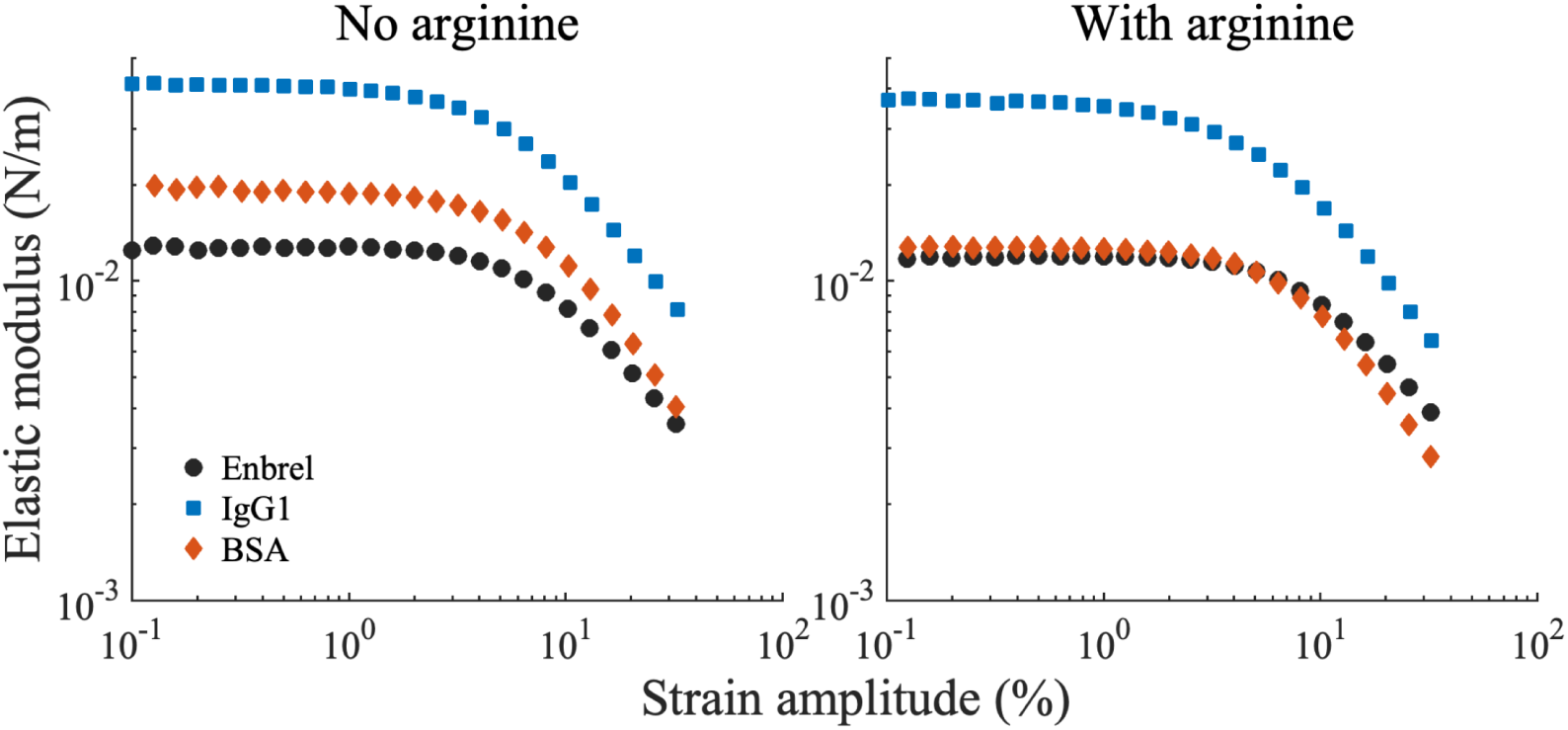
Variations in the elastic modulus as a function of strain amplitude for Enbrel, IgG1, and BSA at a concentration of 0.1 mg/mL. Two plots represent measurements in the absence and presence of arginine.

Figure 1 presents the results for the three proteins (Enbrel, IgG1, and BSA), both in the presence and absence of arginine. It can be observed that the elasticity of IgG1 is higher compared to the other proteins. Additionally, results for the protein solutions with arginine shows that the relative order of the elasticity values remains the same as in the solutions without arginine, indicating that IgG1 molecules form stronger networks at the interface compared to Enbrel and BSA. While arginine does affect the elastic modulus to some extent, it is evident that the forces between the protein molecules remain the dominant factor in determining the strength of the network.

Another important point that can be derived from this figure is the position of the yielding point for different samples. At the yielding point, the protein network starts to be disrupted. The strain amplitude at which this phenomenon occurs is called the critical strain (*γ_c_*), and the associated stress is called the critical stress (*τ_c_*). The critical strain (*γ_c_*) measures the deformability of the network and indicates the extent to which the molecules can be deformed without disintegration [42]. Figure 1 shows that although the proteins exhibit different elasticity in the linear regime, the approximate value of the critical strain remains similar for all three proteins, both in the presence and absence of arginine.

The variations in viscoelasticity of BSA, Enbrel, and IgG1 with frequency are shown in Figure 2. The protein concentration was kept constant at 0.01 mg/mL and the strain amplitude was 2%. The figure shows that for all proteins, the elastic modulus is higher than the loss modulus, both in the presence and absence of arginine, indicating a gel-like network at the interface. Additionally, the evolution of the viscoelastic modulus shows that increasing the frequency leads to an increase in both the elastic and loss moduli. It is important to note that this trend signifies the linear viscoelastic (LVE) regime. In the non-linear regime, a decrease in viscoelasticity is expected following the persistent increase in the linear regime. Figure 2 also illustrates the effects of arginine on the viscoelasticity of the protein layer at the interface. Notably, arginine decreases the viscoelastic moduli of Enbrel and IgG1 while it increases these parameters for BSA at the specified concentration. This effect highlights the fact that the impact of arginine varies depending on its concentration and the intrinsic properties of the proteins, as reflected in our results.

**Figure 2:**
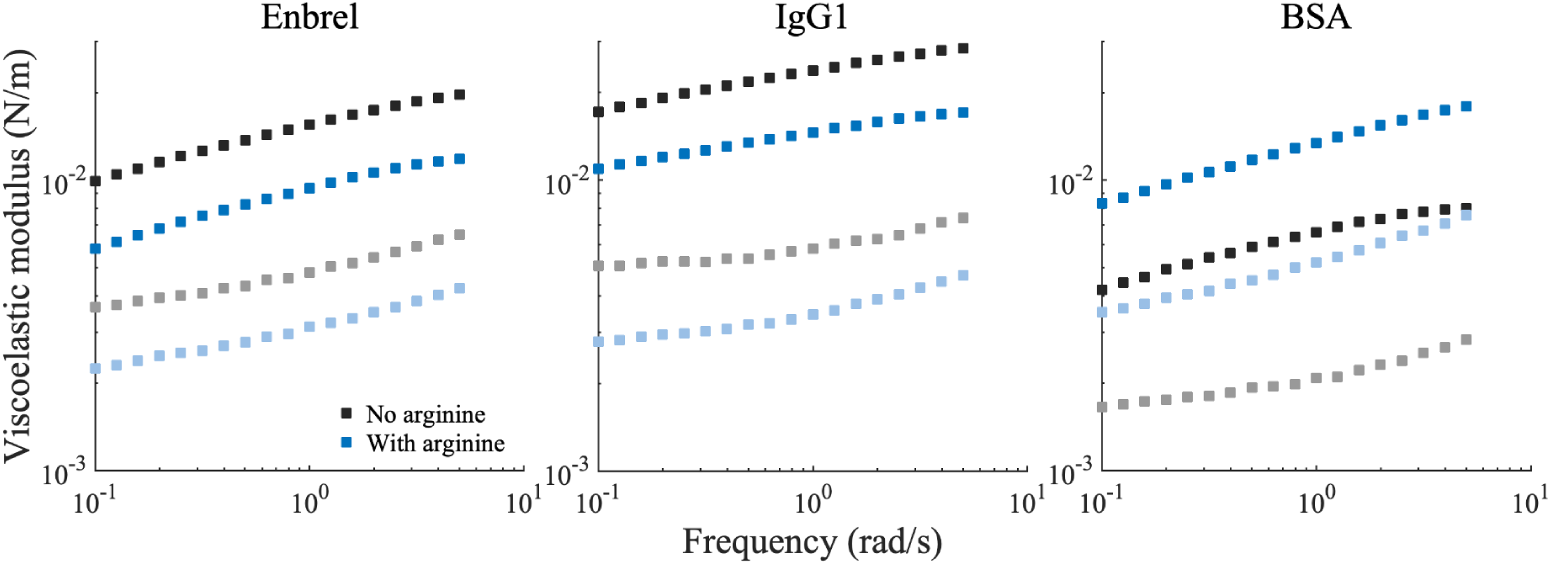
Variations of the elastic modulus (*G*′, symbols with dark color) and loss modulus (*G*′′, symbols with light color) as a function of frequency for Enbrel, IgG1, and BSA at a concentration of 0.01 mg/mL. Each plot shows two conditions: without arginine and with 25 mM arginine added to the solution.

The time-dependent variations in the elastic and loss moduli for the three proteins at different concentrations are shown in Figure 3. A comparative analysis of these plots reveals distinct behaviors among the proteins investigated in this study. The two plots corresponding to Enbrel in Figures 3 present the data for Enbrel at four different concentrations. It is evident that the lowest concentration (0.001 mg/mL) exhibits significantly weaker viscoelastic properties compared to the other three concentrations. The solution with 0.001 mg/mL Enbrel demonstrates negligible elasticity for approximately one hour. Following this period, initial non-zero values for elasticity are observed, which subsequently increase over time. Similar trends are observed in the loss modulus variations. Nonetheless, during the initial stage, when the solution shows no elasticity, non-zero values for the loss modulus are measured, signifying an initial predominance of viscous properties. This trend is similarly observed for other concentrations, with the notable difference that non-zero values for elasticity are achieved at much earlier times at higher concentrations, which indicates a more rapid formation of molecular networks. Furthermore, both plots for Enbrel in Figure 3 show a decline in the rate of increase for both the elastic and loss moduli after a certain time. However, this transition appears to occur later for the lowest concentration (0.001 mg/mL), compared to higher concentrations. It can also be inferred from Figures 3(a-b) that both elastic and loss moduli for the concentrations of 0.01, 0.1, and 1 mg/mL are higher compared to those for the solution of 0.001 mg/mL Enbrel. However, the measured values for these concentrations, particularly over longer time periods, are very similar. To quantify this observation, we extracted the values of the elastic (G^′^) and loss (G^′′^) moduli at an aging time of t=20,000 s, and found that the average coefficient of variation across the three concentrations was only 5.5%. Furthermore, it is noteworthy that during the initial stages, that Enbrel at 1 mg/mL exhibits lower viscoelasticity compared to 0.01 and 0.1 mg/mL.

**Figure 3:**
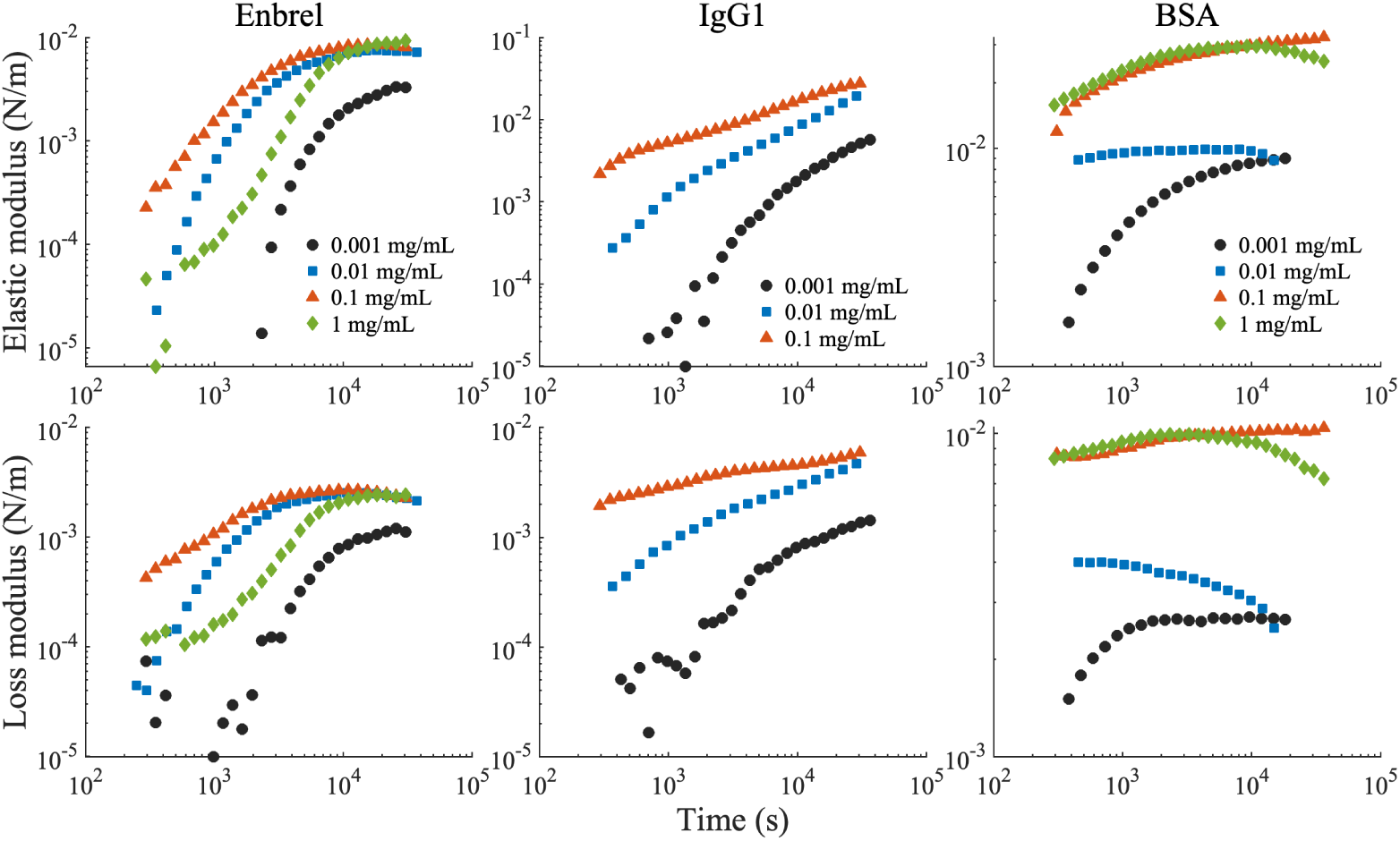
Time evolution of the elastic and loss moduli for the solutions of Enbrel, IgG1, and BSA without arginine. The frequency and strain amplitude were kept constant at 1 rad/s and 2%, respectively.

The data presented for IgG1 in Figure 3 indicates distinct behavior of this protein compared to Enbrel. It should be noted that the concentrations of 0.001, 0.01, and 0.1 mg/mL are considered for IgG1. According to the plots in the middle column of Figure 3, the viscoelasticity of IgG1 solutions at all concentrations continues to increase with an almost constant slope, in contrast to Enbrel solutions, particularly at higher concentrations, which exhibit a decrease over prolonged periods. Furthermore, as the concentration increases, both the elastic and loss moduli increase, although the values for the concentrations of 0.01 and 0.1 mg/mL converge as time approaches 10 hours.

Considering the variations in viscoelastic properties of BSA (third column in Figure 3), distinct characteristics are observed when compared to Enbrel and IgG1. As illustrated in Figure 3, BSA at 0.001 mg/mL, forms a stronger interfacial molecular network and reaches equilibrium faster than Enbrel and IgG1 at the same concentration. Even at higher concentrations, BSA exhibits an earlier onset of the plateau in viscoelasticity relative to the other proteins. This trend is consistent with the findings of Sharma et al. [43], who reported that equilibrium in the interfacial viscosity of BSA was typically reached within 1000 seconds across various concentrations. In our study, however, we additionally observe that at certain BSA concentrations (0.01 and 1 mg/mL), viscoelastic moduli initially plateau but subsequently decrease over time; a behavior not detected in Enbrel or IgG1 at any tested concentration.

The measured elastic moduli for BSA in our experiments range from approximately 0.009 N/m at 0.001 mg/mL to 0.0279 N/m at 1 mg/mL (measured at *t* = 20,000 s), which align well with previously reported values in the literature. For instance, Jaishankar et al.[44] reported G^′^ in the range of 0.021–0.022 N/m for BSA at 50 mg/mL, while Burgess and Sahin[45] reported values near 0.0123 N/m for BSA at 1 mg/mL. Although differences in buffer composition and protein concentration exist between studies, the magnitude of viscoelasticity observed in our experiments is of comparable order.

The impact of arginine on the time-dependent surface properties of the studied proteins is illustrated in Figure 4. Enbrel solutions follow a similar trend to those without arginine (Figure 3), where viscoelasticity increases with protein concentration from 0.001 to 0.1 mg/mL. However, at 1 mg/mL, the equilibrium viscoelasticity is lower.

**Figure 4:**
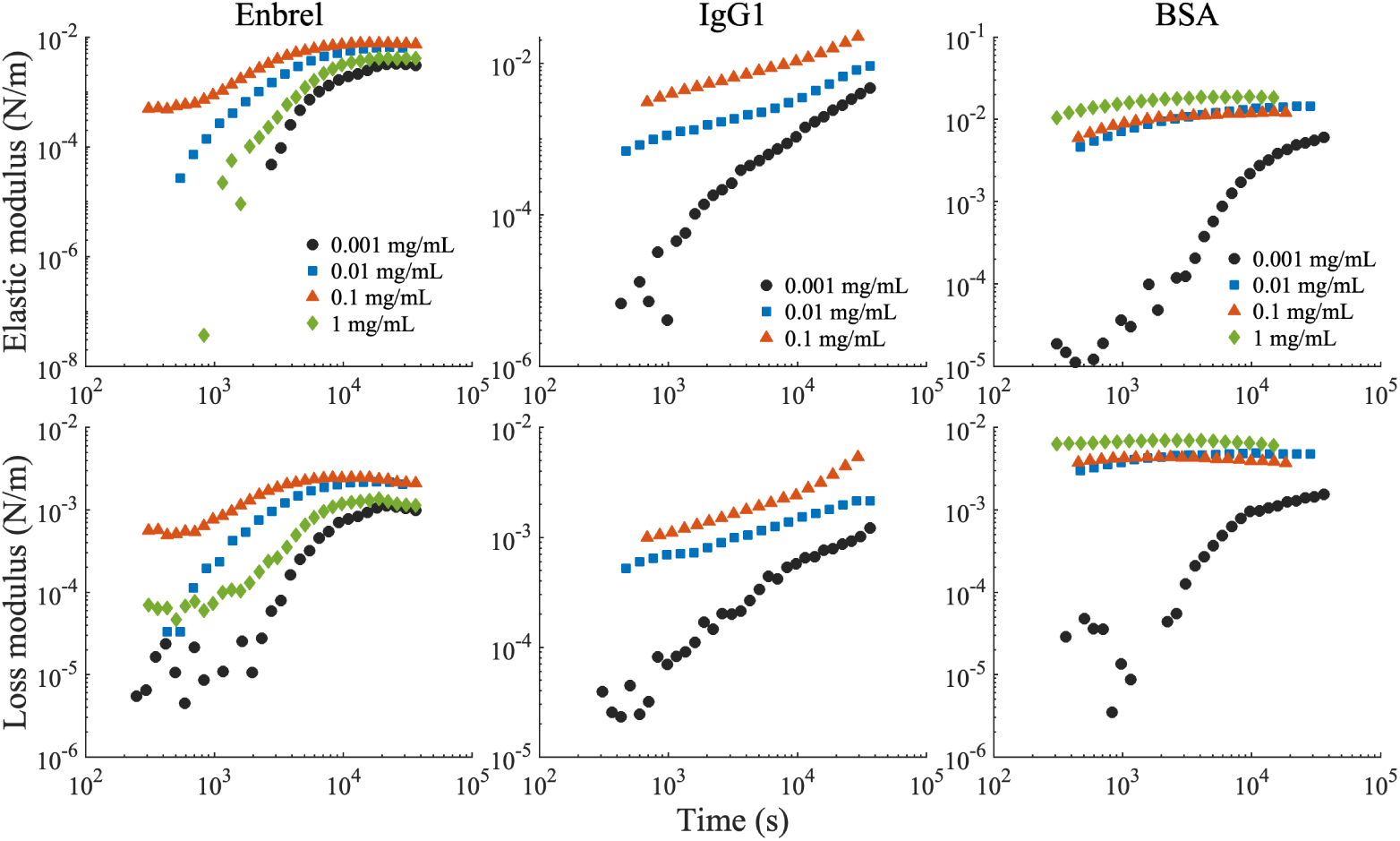
Variations of the elastic and loss moduli in terms of time for the solutions of Enbrel, IgG1, and BSA that contain 25 mM arginine. The frequency and strain amplitude were kept constant at 1 rad/s and 2%, respectively.

Figure 4 also illustrates the behavior of IgG1 in the presence of arginine. As can be seen, increasing the IgG1 concentration results in a rise in the viscoelastic moduli of the protein solutions. Similar to the condition where arginine was absent (Figure 3), it is evident that the slope of the viscoelastic moduli increases for Enbrel (first column of Figure 4 diminishes after a certain period. However, the viscoelasticity of IgG1 solutions continues to increase at a steep rate.

Based on our observations of monoclonal antibodies, the samples containing BSA and arginine exhibit distinct behaviors at the air-water interface. Figure 4 demonstrate that BSA at a concentration of 0.001 mg/mL has significantly lower elastic and loss moduli. As the concentration increases, a sudden increase in viscoelasticity is observed; however, the values for the concentrations of 0.01, 0.1, and 1 mg/mL are close, with only slight increases in viscoelasticity across this range. As an estimate, the average coefficient of variation for the elastic and loss moduli at t = 20,000 s is 21%, indicating that the viscoelastic response does not significantly change as the concentration increases from 0.01 to 1 mg/mL.

In contrast to IgG1, the viscoelasticity increment rate for BSA at the lowest concentration decreases over time but does not reach equilibrium within the time frame considered. At higher concentrations, however, equilibrium is reached much more quickly. Additionally, when comparing these plots to those in Figure 3, it is clear that the elastic and loss moduli of BSA solutions with arginine do not decrease over time, unlike the solutions without arginine.

Comparing the results presented in Figures 3, we observe that, in general, the interfacial viscoelasticity of the protein solutions decreases upon the addition of arginine. To better quantify this trend, we calculated the percentage change in the elastic (G^′^) and loss (G^′′^) moduli at *t* = 20,000 s for each protein as arginine was added at each concentration. Our findings show that the average percentage change—computed across all concentrations and for both G^′^ and G^′′^—was *−*16.4% for Enbrel, *−*37.7% for IgG1, and *−*14.5% for BSA.

Moreover, it is important to note that similar tests conducted with the buffer solution containing arginine (PBS + arginine) showed that the viscoelasticity remained close to zero at all times. Consequently, it can be deduced that network formation at the interface is primarily influenced by protein molecules, although arginine may alter their interactions and connections with each other.

The data obtained from relaxation tests under 2% step strain for the three proteins is presented in Figure 5. All solutions have a concentration of 0.001 mg/mL, and the figure illustrates both the cases in presence and absence of arginine. As shown in Figure 5, the interface of the IgG1 sample exhibits a longer relaxation time, and the stress measured for this solution is higher compared to the other proteins. Conversely, the Enbrel solution relaxes more rapidly, which indicates weaker intermolecular interactions. In the scenario where arginine is added to the solution, as depicted in Figure 5, Enbrel for this specific concentration shows similar behavior to that without arginine. Another important observation is that the slope of stress reduction, or relaxation, for Enbrel and IgG1 is similar in both plots, while BSA relaxes with a steeper slope in the presence of arginine.

**Figure 5:**
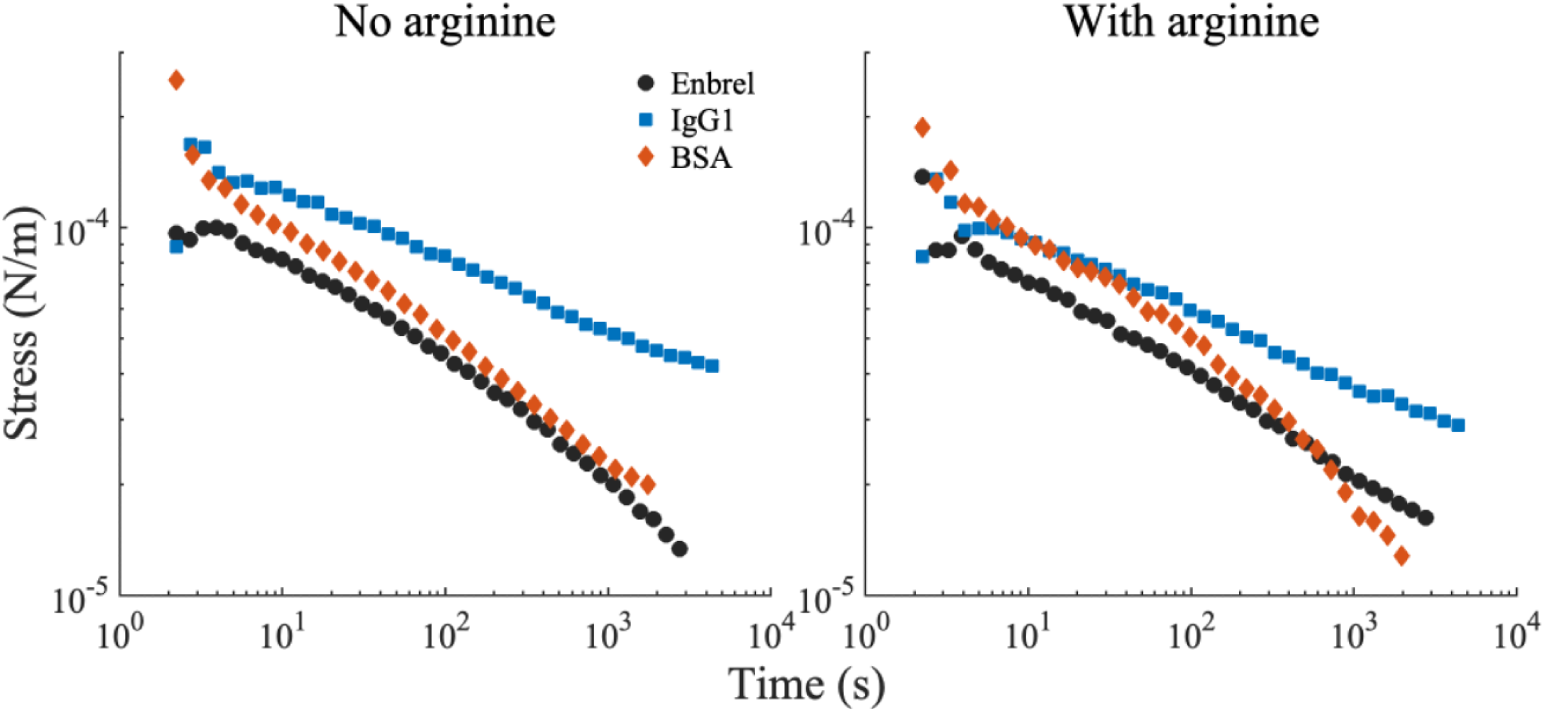
Stress relaxation at 2% strain for Enbrel, IgG1, and BSA at a concentration of 0.001 mg/mL.

Furthermore, the viscosity of Enbrel and IgG1 solutions at varying shear rates was measured, with data obtained for a protein concentration of 0.1 mg/mL, both in the absence and presence of arginine, as shown in Figure 6. All solutions exhibit shear-thinning behavior under shear deformation, indicating the disruption of molecular networks at the air-water interface as the shear rate increases. A key distinction between Enbrel and IgG1 is that Enbrel demonstrates yielding behavior at low shear rates. Additionally, compared to the arginine-containing solution, IgG1 without arginine exhibits a slightly higher viscosity at low shear rates. However, at higher shear rates, the viscosity of both Enbrel and IgG1 becomes nearly identical, irrespective of arginine presence.

**Figure 6:**
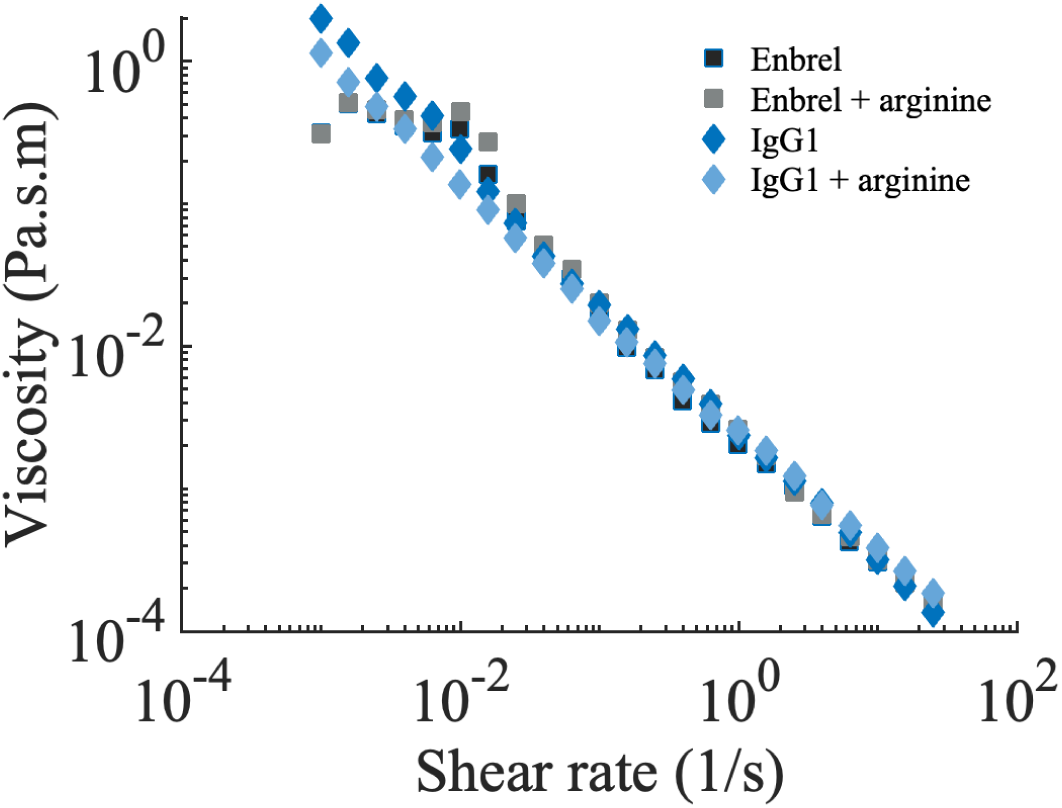
Plot of shear viscosity versus shear rate for Enbrel and IgG1 solutions, comparing conditions with and without the presence of arginine.

The behavior of proteins at interfaces can significantly differ under extreme shear or strain compared to their behavior in the linear viscoelastic regime, as observed in the results presented earlier in this paper. We aimed to investigate how protein surfaces behave at large strains and the influence of arginine on this behavior. To do so, we measured the stress response of solution interfaces subjected to a single cycle of oscillatory deformation at a constant frequency of 1 rad/s, comparing six different samples at a strain amplitude of 100%. It is important to note that our objective was to gain a general understanding of how the nonlinear response of these solution interfaces differs from their linear rheology. Therefore, we examined only one concentration and a single strain amplitude for each protein. Further studies would be valuable for a more detailed characterization of the nonlinear behavior of these three proteins.

Figure 7 presents the elastic Lissajous curves of stress versus strain for the highest concentration studied of each protein (Enbrel and BSA at 1 mg/mL, and IgG1 at 0.1 mg/mL), both with and without arginine. The corresponding viscous Lissajous curves of stress versus strain rate are shown in Figure 8. Figure 7 illustrates how the curve shapes vary depending on protein type and the presence of arginine. The first key observation is that all three protein solutions—Enbrel, IgG1, and BSA—exhibit viscoelastic behavior. This is evident from the elliptical or looped shapes in both the elastic (Figure 7) and viscous (Figure 8) curves. The deviation from a perfectly elliptical shape indicates a combination of elastic and viscous properties, where energy is both stored during deformation and dissipated as heat.

**Figure 7:**
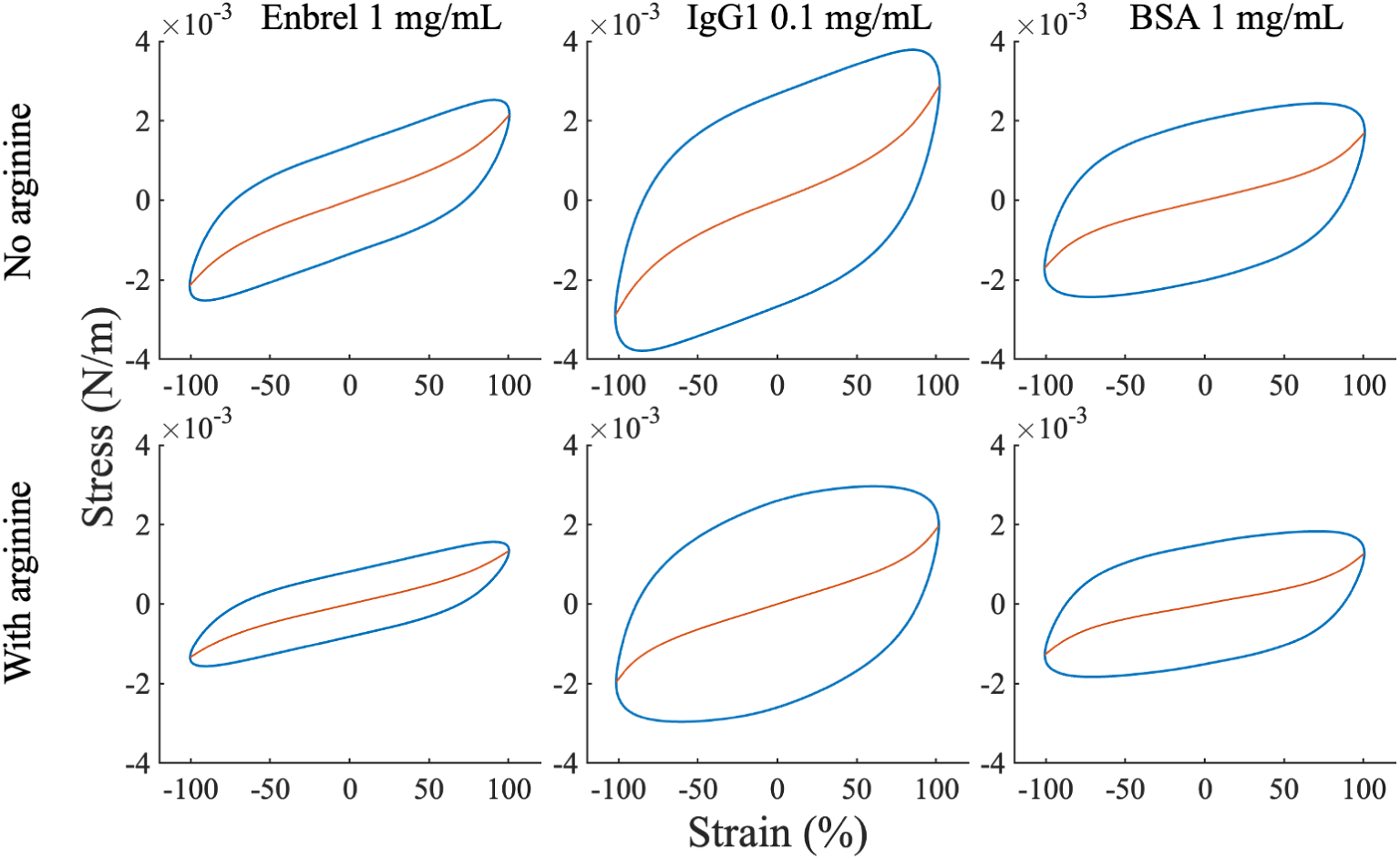
Elastic Lissajous–Bowditch curves obtained from large amplitude oscillatory shear (LAOS) tests for Enbrel, IgG1, and BSA at 0.1 mg/mL, both with and without 25 mM arginine. The blue lines depict the total stress, while the red lines represent the elastic stress.

**Figure 8:**
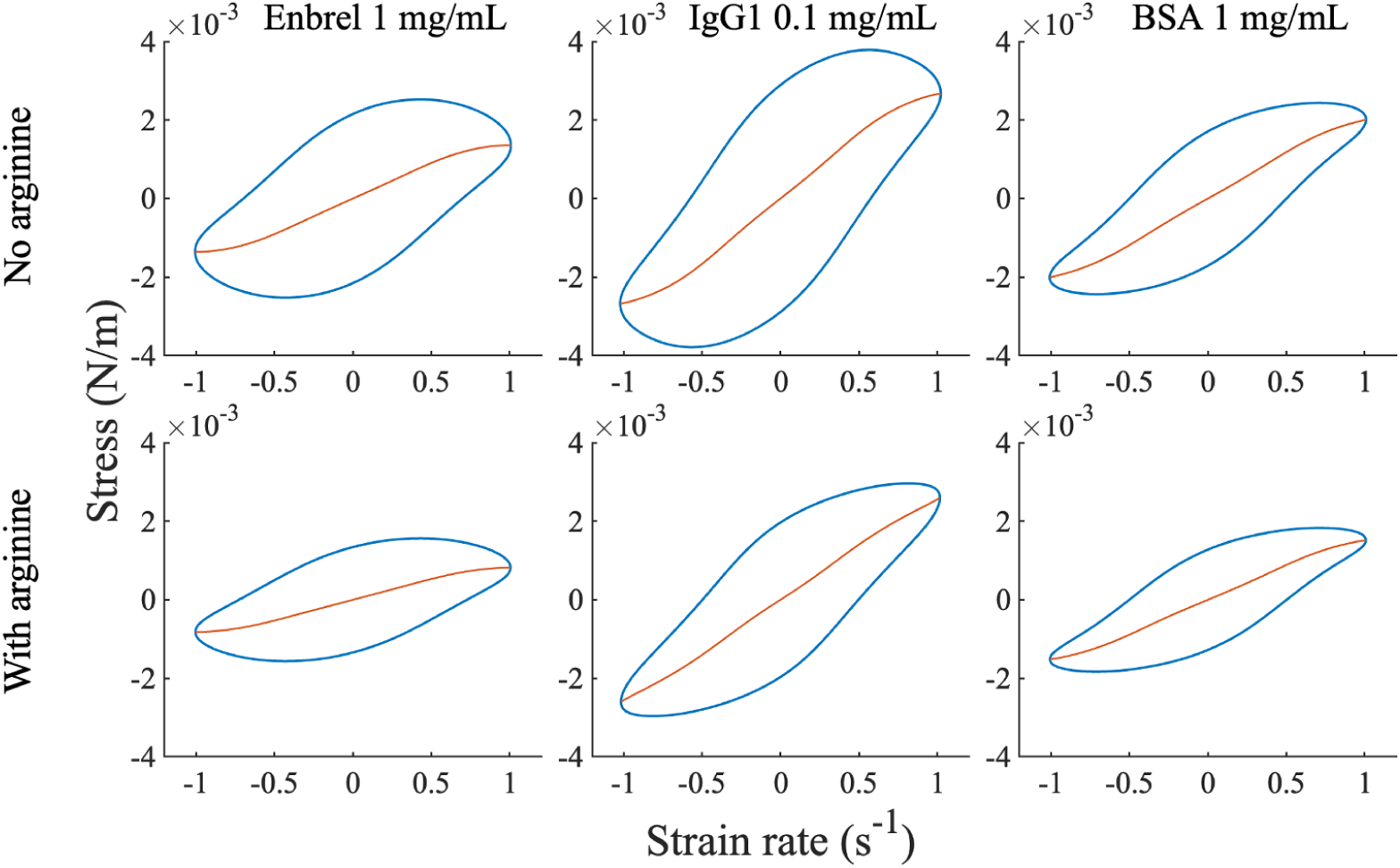
Viscous Lissajous–Bowditch curves obtained from large amplitude oscillatory shear (LAOS) tests for Enbrel, IgG1, and BSA at 0.1 mg/mL, both with and without 25 mM arginine. The blue lines depict the total stress, while the red lines represent the viscous stress.

Furthermore, the viscoelastic properties of the three proteins differ under the tested conditions. IgG1’s profile shows more distortion from an elliptical shape, and greater intra-cycle nonlinearity with pronounced tip bending at large strains, whereas Enbrel exhibits a tighter loop with smaller hysteresis area, indicating reduced viscous dissipation at *γ*_0_ = 100% and the tested *ω* (Figure 7). BSA shows intra-cycle viscoelastic properties that fall between those of Enbrel and IgG1. These differences suggest differences in interfacial microstructure, protein–protein interaction strength, or the relaxation-time spectrum of the protein layers at the tested concentrations.

It is well-established that the area enclosed by the elastic Lissajous curves indicates the viscous dissipation within the system [38, 46, 47]. Based on this, arginine appears to systematically influence the viscoelastic behavior of all three proteins. Specifically, its addition reduces the intra-cycle viscous dissipation. This is evidenced by the decreased enclosed areas in the elastic Lissajous curves for each protein by adding arginine (Figure 8).

The strain-stiffening (S) and shear-thickening (T) ratios for the protein samples discussed in Figures 7 and 8 are presented in Table 1. As S and T deviate further from zero, the intra-cycle nonlinearity of the protein interface becomes more pronounced. All protein formulations exhibit positive T and negative S values, indicating intra-cycle strain-stiffening and shear-thinning behavior. However, the magnitudes of these parameters vary depending on protein type and the presence of arginine, likely due to differences in intermolecular interactions. Specifically, for IgG1 and BSA, arginine reduces T, suggesting a lower degree of strain-stiffening, whereas Enbrel shows negligible changes in T upon arginine addition. Additionally, arginine decreases the absolute value of S, though this effect is minimal for BSA. Among the three proteins, Enbrel exhibits the highest absolute S value, indicating the most pronounced shear-thinning behavior.

**Table 1:**
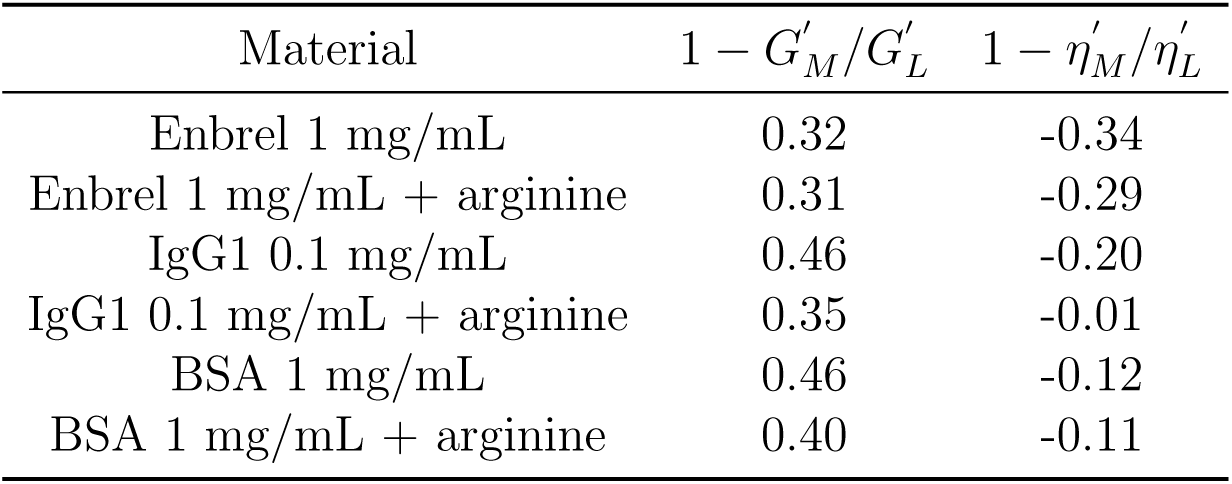
Strain-stiffening and shear-thickening ratios for various protein formulations.

### 3.2. ζ-potential

Protein-protein interactions are influenced by various factors, including hydrophobic interactions, dipole-dipole interactions, and electrostatic forces [24]. The electrostatic forces can be one of the key factors that impact the interfacial rheological properties of proteins. To gain a better understanding of the role of electrostatic interactions, we measured the *ζ*-potential values of all three proteins at a concentration of 0.1 mg/mL, both with and without the addition of arginine. Additionally, we assessed the *ζ*-potential for Enbrel at a higher concentration of 1 mg/mL to evaluate the impact of protein concentration on this parameter. The obtained results are presented in Table 2.

**Table 2:** ζ-potential measurement results for various protein formulations.

| Materials | ζ-potential(mV) | |
| --- | --- | --- |
|  | Without Arginine | With Arginine |
| Enbrel 1 mg/mL | $-6.09 \pm 0.64$ | $-4.48 \pm 0.12$ |
| Enbrel 0.1 mg/mL | $-6.15 \pm 0.76$ | $-4.31 \pm 0.22$ |
| IgG1 0.1 mg/mL | $0.91 \pm 0.19$ | $-3.5 \pm 0.32$ |
| BSA 0.1 mg/mL | $-7.22 \pm 0.54$ | $-6 \pm 0.72$ |

Examining the information in Table 2, one can see that the Enbrel solution without arginine exhibits a negative *ζ*-potential. Also, a comparison between the concentrations of 0.1 mg/mL and 1 mg/mL reveals that Enbrel’s average charge remains unchanged with an increase in protein concentration. On the other hand, the *ζ*-potential of IgG1 in the absence of arginine is positive. Given that the isoelectric point of IgG1 is above 6.5 and near 8.5 [48, 49], the measured value aligns with the intrinsic properties of this protein. BSA, however, displays a greater absolute charge compared to the other two proteins.

The addition of arginine significantly affects the *ζ*-potential values, as shown in Table 2. For Enbrel and BSA, the absolute *ζ*-potential decreases upon arginine addition. In contrast, IgG1 undergoes a charge reversal from positive to negative, with an increase in its absolute *ζ*-potential in the presence of arginine. Notably, the *ζ*-potential of Enbrel remains relatively unchanged across different concentrations. Furthermore, similar to the arginine-free formulations, BSA with arginine retains the highest absolute *ζ*-potential among the tested proteins.

## 4. Discussion

The kinetics of protein adsorption at the air-water interface and subsequent molecular network formation are influenced by molecular-level factors such as protein structure, molecular size, conformational state, and intermolecular interactions. Upon dissolution in aqueous solutions, proteins begin to adsorb at the interface, followed by the formation of molecular networks. Time-dependent rheological data, illustrated in Figures 3 and 4, show that viscoelastic moduli are initially low but increase over time. This observation indicates that proteins or arginine molecules adsorb at the interface over time and form molecular networks. This phenomenon is time-dependent and varies among different samples due to differences in protein type, concentration, or the presence of arginine as an excipient.

The distinct mechanistic properties of Enbrel and IgG1 highlight the influence of the Fab region in intermolecular interactions. As mentioned in the introduction, Enbrel shares an Fc domain with IgG1 but differs in its Fab region, where tumor necrosis factor-*α* (TNF*α*) is fused to the IgG1 Fc domain. Figures 3 and 4 show that the interfacial viscoelasticity of Enbrel solutions increases initially but then exhibits a reduction in its rate of increase, eventually reaching a plateau. In contrast, IgG1 solutions maintain a high rate of increase even after 10 hours.

The duration of the time-sweep tests extends well beyond the adsorption period of IgG1, as reported by Kannan et al. [50]. We therefore attribute the continued increase in interfacial moduli to post-adsorption processes, namely molecular rearrangements, conformational transitions, or intermolecular associations occurring after adsorption. A noteworthy observation in Figures 3 and 4 is that, particularly at longer aging times, IgG1 exhibits a steeper increase in both elastic and loss moduli than Enbrel. Prior studies [51, 52, 53] have linked the interfacial mechanical response of adsorbed protein layers to molecular interactions driven by post-adsorption conformational rearrangements and unfolding. Consistent with this framework, our data indicate that IgG1 molecules undergo more pronounced conformational rearrangement and over longer timescales, which result in higher moduli and growth rates. Accordingly, IgG1 appears more surface-active and more prone to interfacial aggregation than Enbrel. This difference arises from the Fab region’s distinct structural and physicochemical properties, potentially due to increased hydrophobicity, electrostatic interactions, or other molecular forces that promote intermolecular associations.

BSA exhibits inherent structural and conformational differences from antibodies, which are reflected in the rheological data shown in Figures 3 and 4. A key distinction between BSA and the other two proteins (Enbrel and IgG1) is that, particularly at higher concentrations (0.01, 0.1, and 1 mg/mL), BSA at the initial time points displays a greater elastic modulus than the corresponding solutions of Enbrel and IgG1. This suggests that for BSA, molecular interactions are stronger and form faster than IgG1, and Enbrel, as as evident in Figures 3 and 4. This behavior can be attributed to BSA’s smaller size, which enables faster adsorption and denser packing at the interface. Additionally, easier conformational changes and unfolding of BSA molecules at the interface could be factors that give rise to protein cross-linking and aggregation, and consequently, higher viscoelasticity [54, 55, 53].

Furthermore, as shown in Figure 4, arginine appears to facilitate BSA unfolding and aggregate formation at the interface, as indicated by the similar viscoelastic responses observed across concentrations of 0.01, 0.1, and 1 mg/mL within the experimental time range. This concentration-independent behavior may suggest that the interface becomes saturated due to extensive unfolding and adsorption of protein molecules. Assuming this scenario—which we consider the most plausible—arginine likely promotes structural rearrangement and unfolding at the interface, thereby maximizing the interfacial area occupied by each protein molecule. As discussed in the Results section, the interfacial viscoelasticity of the buffer formulation containing arginine (PBS+arginine) was negligible, supporting the conclusion that arginine’s impact on interfacial mechanical properties arises primarily through its interactions with protein molecules. These arginine–protein interactions likely modulate protein–protein associations at the interface, rather than contributing via the formation of mechanically active arginine networks. The minimal variation in viscoelasticity across concentrations further supports the idea that arginine modulates protein–protein interactions in a manner that is relatively insensitive to protein concentration. However, when comparing the formulations with and without arginine (Figures 3 and 4), both the elastic and viscous moduli are consistently lower in the presence of arginine for most concentrations. This observation suggests that although arginine enhances interfacial unfolding, it weakens intermolecular interactions and reduces the ability of BSA molecules to form a cohesive viscoelastic network at the interface.

The *ζ*-potential values of proteins provide insight into the extent to which electrostatic forces influence their interfacial behavior. As shown in Table 2, Enbrel exhibits a negative *ζ*-potential both in the presence and absence of arginine. However, the addition of arginine reduces the absolute value of this parameter, indicating a decrease in electrostatic repulsion. A similar trend is observed for BSA. In the case of IgG1, the addition of arginine reverses the charge and causes the *ζ*-potential to switch from positive to negative.

Comparison of the *ζ*-potential measurements with the interfacial rheology results for selected protein formulations indicates that the absolute value of the *ζ*-potential alone does not dictate interfacial elasticity, and by extension, molecular network strength. For instance, Enbrel with arginine exhibits a lower absolute *ζ*-potential than Enbrel without arginine, suggesting a reduced electrostatic repulsive barrier for protein–protein interactions. Nevertheless, the addition of arginine results in lower elasticity, as shown in Figures 1, 3, and 4. A second example arises from comparing BSA and Enbrel in the presence of arginine. Table 2 shows that BSA with arginine has a higher absolute *ζ*-potential than Enbrel with arginine. Yet, at a concentration of 1 mg/mL, Figure 4 demonstrates that BSA with arginine achieves higher surface elasticity both initially (indicating faster network formation) and at equilibrium, where its final elasticity also exceeds that of Enbrel with arginine. These results suggest that although electrostatic forces influence interfacial protein interactions, particularly in the presence of arginine, they are not the dominant factor. Instead, the observed differences in the interfacial rheology of arginine-containing formulations are more likely governed by hydrophobic interactions [56]. It should be noted that the *ζ*-potential values reported here correspond to bulk solution measurements. However, protein adsorption, conformational rearrangement, and partial unfolding at the interface may expose different molecular regions that can potentially alter surface charge. This possible phenomenon is not considered in this analysis. Nonlinear mechanical response of protein formulations uncovers additional distinctions in their interfacial properties. As shown in Figure 7, IgG1 solutions exhibit greater nonlinearity, higher viscous dissipation, and a higher maximum stress response compared to Enbrel solutions. Additionally, Table 1 indicates that IgG1 samples demonstrate a more pronounced strain-stiffening behavior and less shear-thinning, which suggests greater resistance to deformation. This observation is consistent with the trends observed in the linear rheology results (Figures 3, 4, and 5). Furthermore, the addition of arginine to Enbrel appears to reduce the maximum stress response at the interface and decrease the area enclosed by the Lissajous–Bowditch curves, indicating lower viscous dissipation (Figure 7). However, Table 1 suggests that the strain-stiffening ratio of Enbrel remains almost unchanged upon arginine addition, while its shear-thinning properties decrease. It is important to note that the trends observed in Figures 7 and 8 pertain to the intra-cycle response of the interface. For instance, Figure 1 demonstrates that protein solutions exhibit strain-softening behavior as strain amplitude increases, which reflects inter-cycle behavior. In contrast, the intra-cycle behavior of the interface under strain follows an opposite trend (Table 1). Nevertheless, in both cases, the observed variations in rheological properties due to formulation differences provide valuable insights into the interfacial dynamics of these protein solutions.

Another noteworthy observation is the time evolution of Enbrel’s viscoelasticity at a concentration of 1 mg/mL compared to lower concentrations. Figure 3 shows that the viscoelasticity of the Enbrel interface initially increases as the concentration rises from 0.001 to 0.1 mg/mL but decreases when the concentration increases further to 1 mg/mL. A similar trend is observed for Enbrel in the presence of arginine (Figure 4). This behavior may be attributed to the adsorption rate of Enbrel molecules at the interface. At higher concentrations, the reduced intermolecular distance may enhance repulsive interactions, thereby lowering the adsorption rate. However, at equilibrium (Figure 3), Enbrel’s viscoelasticity varies little across concentrations from 0.01 to 1 mg/mL.

Arginine is known to function both as a salt and as a molecule capable of forming hydrogen bonds and hydrophobic interactions with proteins [12]. These properties become particularly important at the air-water interface, where protein molecules undergo complex rearrangement and conformational changes, which can potentially expose their hydrophobic regions [57, 58]. Consequently, arginine can interact with proteins in multiple ways, either promoting or preventing protein aggregation depending on various factors, such as protein abundance at the interface, protein type, pH, the presence of other excipients, and more.

Our results indicate that the effects of arginine vary. For BSA and Enbrel, arginine can either strengthen molecular interactions (Figures S1 and S3 in the Supplementary material) or weaken them, though in most formulations, we observe a reduction in viscoelasticity. For IgG1 (Figure S2 in the Supplementary material), arginine appears to weaken molecular interactions, although this effect is negligible at a concentration of 0.1 mg/mL.

This work highlights the importance of evaluating interfacial behavior when selecting excipients for therapeutic protein formulations. While arginine is well known for suppressing protein–protein interactions in solution, our interfacial rheology results demonstrate that it also reduces protein self-association and viscoelastic film formation at the air–water interface. However, despite this reduction, some degree of interfacial aggregation persists, indicating that arginine alone may not fully prevent surface-induced destabilization. This suggests that formulation strategies should not only aim to inhibit aggregation in the bulk phase but also prioritize minimizing residual interfacial interactions that could compromise protein stability over time. The identification of alternative or complementary excipients that provide improved interfacial stability without compromising safety remains a key objective. Furthermore, our work underscores the strength of interfacial rheology as a sensitive analytical tool to probe molecular network formation and protein adsorption behavior. Applying interfacial rheology at early formulation stages can provide critical insights for selecting optimal excipients and designing proteins with reduced susceptibility to interfacial aggregation. Together, these approaches can contribute to the development of safer, more stable, and manufacturable protein therapeutics.

## 5. Conclusion

We evaluated whether arginine mitigates interfacial protein–protein interactions and aggregation of Enbrel (a fusion protein) at the air–water interface by comparing it’s behavior with that for IgG1, and BSA across multiple concentrations, with and without arginine, under linear (small-strain) and nonlinear (large-strain) interfacial deformation. On average, arginine decreased the interfacial viscoelasticity by 16.4% for etanercept, 37.7% for IgG1, and 14.5% for BSA. Despite this attenuation, the absolute moduli (Figure 4) still indicate appreciable adsorption and subsequent network formation in the presence of arginine.

*ζ*-potential measurements showed that arginine reduced the magnitude of the (bulk) *ζ*-potential, consistent with weaker electrostatic repulsion. However, the trends in interfacial viscoelasticity did not mirror the changes in *ζ*-potential, indicating that interfacial mechanics cannot be inferred from bulk electrostatics alone. This interfacial picture differs from prior reports that emphasize predominantly electrostatic arginine–protein interactions in bulk solution [21, 22, 26]. Furthermore, the marked reduction in BSA interfacial viscoelasticity with arginine (Figures 3 and 4) and prior reports of negligible changes in bulk viscosity upon arginine addition [24, 27], support the conclusion that the nature of the protein–amino acid interactions differs at interfaces versus in bulk solution.

In addition, our results demonstrated that all protein formulations exhibited dominant elastic behavior after only a few minutes of aging—consistent with previous reports for BSA [31, 43, 59] and monoclonal antibodies [32, 60]—and that this behavior persisted even upon addition of arginine (Figure 4). Importantly, a novel observation from this study is that time-dependent viscoelasticity measurements (Figures 3 and 4), relaxation tests (Figure 5), and non-linear rheological analyses (Figures 7 and 8) consistently revealed greater interfacial activity for IgG1 compared to Enbrel. This difference is likely rooted in the structural variations between the TNF*α* receptor domain of Enbrel and the Fab domain of IgG1.

The interfacial behavior of proteins across different concentrations revealed that, for certain systems, specifically Enbrel (both with and without arginine) and BSA with arginine, the viscoelastic response exhibited only minimal changes within the concentration range of 0.01 to 1 mg/mL. This outcome differs markedly from earlier reports on the mechanistic behavior of proteins in the bulk solution, where the viscosity keeps rising with concentration, persisting up to very high levels for mAbs [61, 62] and BSA in the presence of arginine [63]. Such differences may arise from saturation of the interface or the establishment of an equilibrium state governing protein conformational dynamics at the interface. Further study is needed to probe broader concentration ranges and to assess if similar interfacial mechanistic behavior is maintained at higher concentrations, particularly in systems where proteins interact with amino acids.

Using the standard LAOS Lissajous-Bowditch procedure [38], we constructed parametric *σ*-*γ* and *σ*-*γ̇* trajectories from the interfacial oscillation data at a strain amplitude of 100%. Analysis of these curves showed greater intra-cycle strain-stiffening and higher peak stress for IgG1 than for Enbrel in the nonlinear regime. This trend is consistent with our linear-regime results and supports stronger interfacial molecular interactions for IgG1. Addition of arginine decreased the area enclosed by the elastic Lissajous-Bowditch loops (energy dissipated per cycle) and reduced the cycle-maximum stress for all three proteins, indicating weakened interfacial network connectivity in the presence of arginine.

Building upon our findings, future research should investigate protein interfacial behavior at a broader range of interfaces, including solid–liquid and oil–water boundaries, which are commonly encountered throughout the biopharmaceutical lifecycle, from manufacturing and storage to administration. Moreover, the inclusion of multiple excipients in commercial formulations introduces additional complexity. Therefore, further investigation is needed to elucidate their combined effects on protein stability and interfacial interactions.

## 6. Acknowledgment

We thank Merck & Co., Inc., Rahway, NJ, USA for their financial support of this study. We also acknowledge Suzette Pabit for her expert review and insightful suggestions that helped improve the manuscript.

