## Supplementary Material for "Effects of arginine on the interfacial behavior of proteins"

Arezoo M. Ardekani^∗,†^

†School of Mechanical Engineering, Purdue University, West Lafayette, IN, United States

of America

‡Merck &Co., Inc., Rahway, N J, 07065

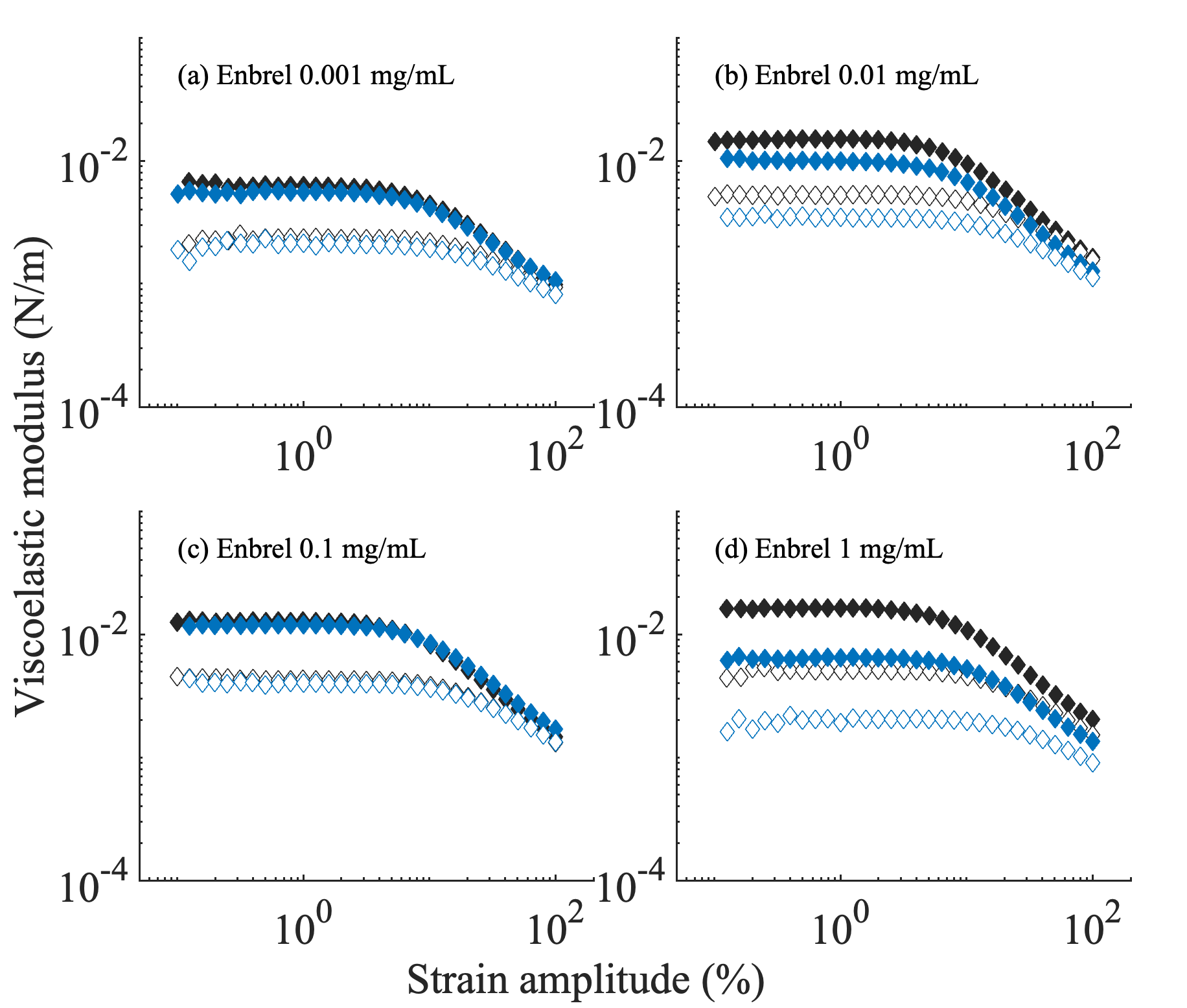

**Figure S-1**: Elastic and loss modulus as a function of strain amplitude for Enbrel at concentrations of (a) 0.001 mg/mL, (b) 0.01 mg/mL, (c) 0.1 mg/mL, and (d) 1 mg/mL. For all concentrations, data without arginine (black markers) and with arginine (blue markers) are shown. Open markers represent the loss modulus, while solid markers correspond to the elastic modulus.

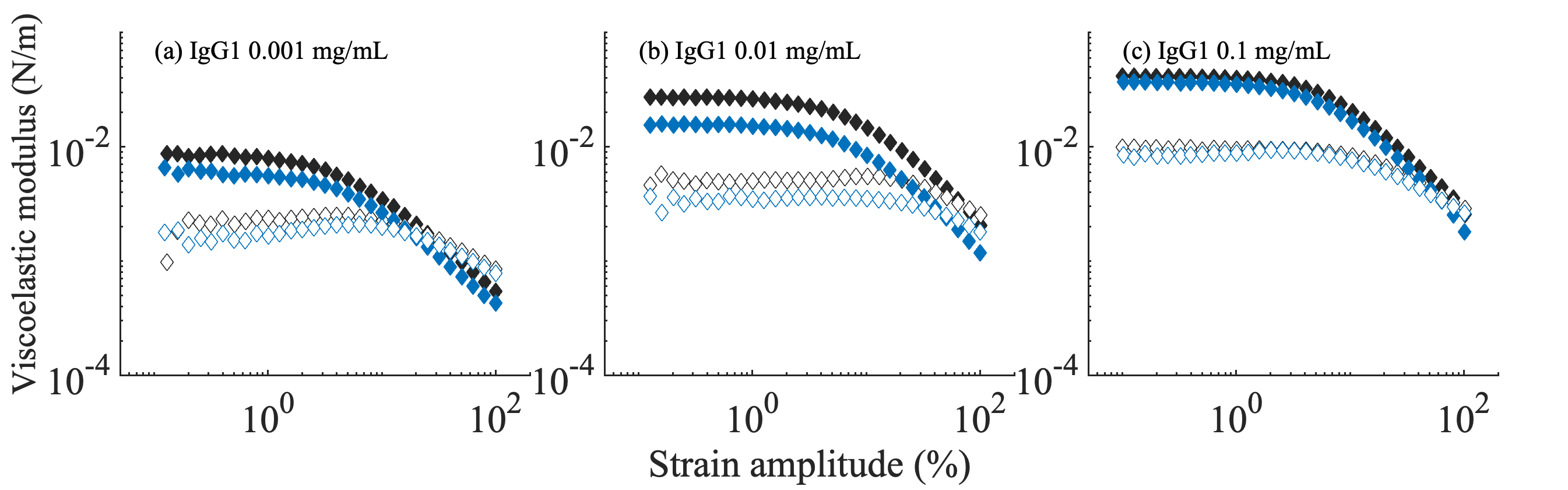

**Figure S-2**: Elastic and loss modulus as a function of strain amplitude for IgG1 at concentrations of (a) 0.001 mg/mL, (b) 0.01 mg/mL, and (c) 0.1 mg/mL. For all concentrations, data without arginine (black markers) and with arginine (blue markers) are shown. Open markers represent the loss modulus, while solid markers correspond to the elastic modulus.

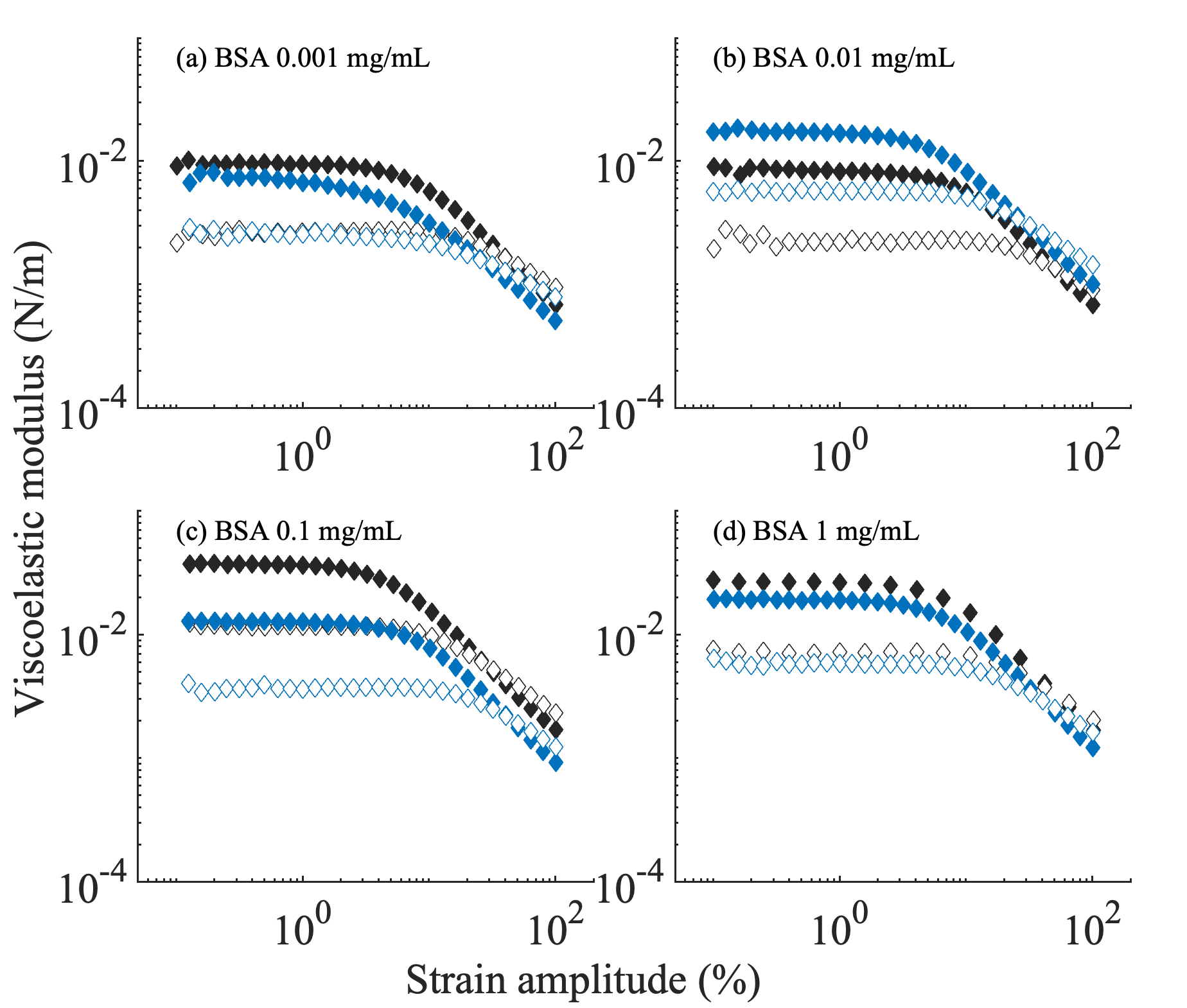

**Figure S-3**: Elastic and loss modulus as a function of strain amplitude for BSA at concentrations of (a) 0.001 mg/mL, (b) 0.01 mg/mL, (c) 0.1 mg/mL, and (d) 1 mg/mL. For all concentrations, data without arginine (black markers) and with arginine (blue markers) are shown. Open markers represent the loss modulus, while solid markers correspond to the elastic modulus.

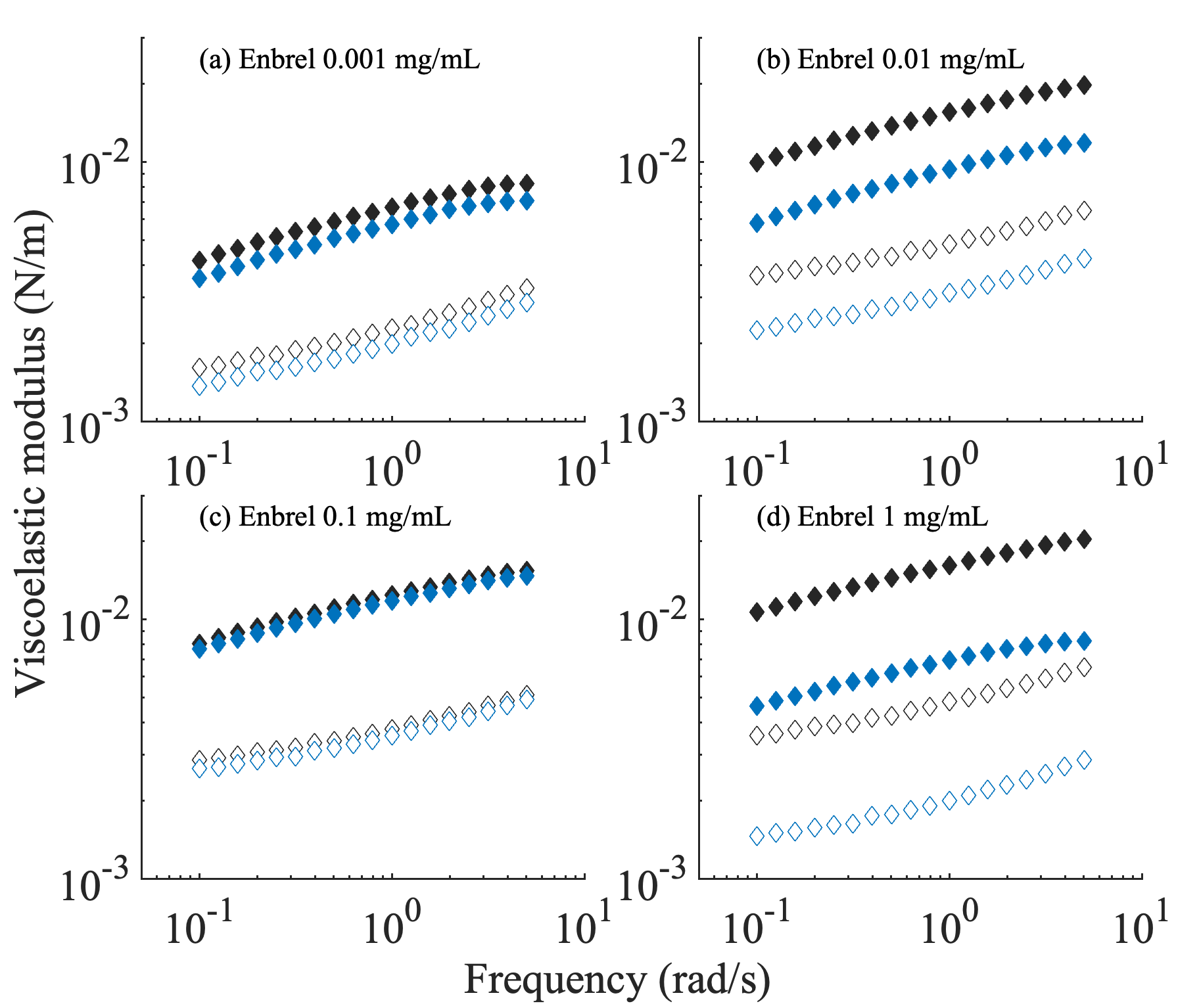

**Figure S-4**: Elastic and loss modulus as a function of angular frequency for Enbrel at concentrations of (a) 0.001 mg/mL, (b) 0.01 mg/mL, (c) 0.1 mg/mL, and (d) 1 mg/mL. For all concentrations, data without arginine (black markers) and with arginine (blue markers) are shown. Open markers represent the loss modulus, while solid markers correspond to the elastic modulus.

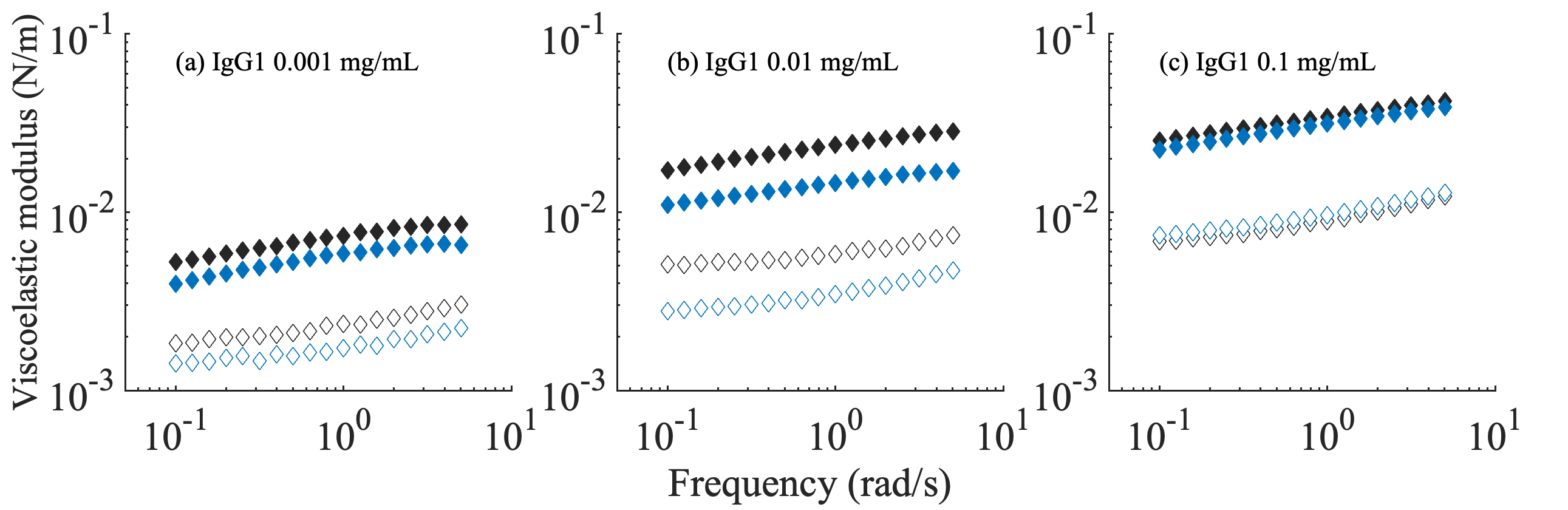

**Figure S-5**: Elastic and loss modulus as a function of angular frequency for IgG1 at concentrations of (a) 0.001 mg/mL, (b) 0.01 mg/mL, and (c) 0.1 mg/mL. For all concentrations, data without arginine (black markers) and with arginine (blue markers) are shown. Open markers represent the loss modulus, while solid markers correspond to the elastic modulus.

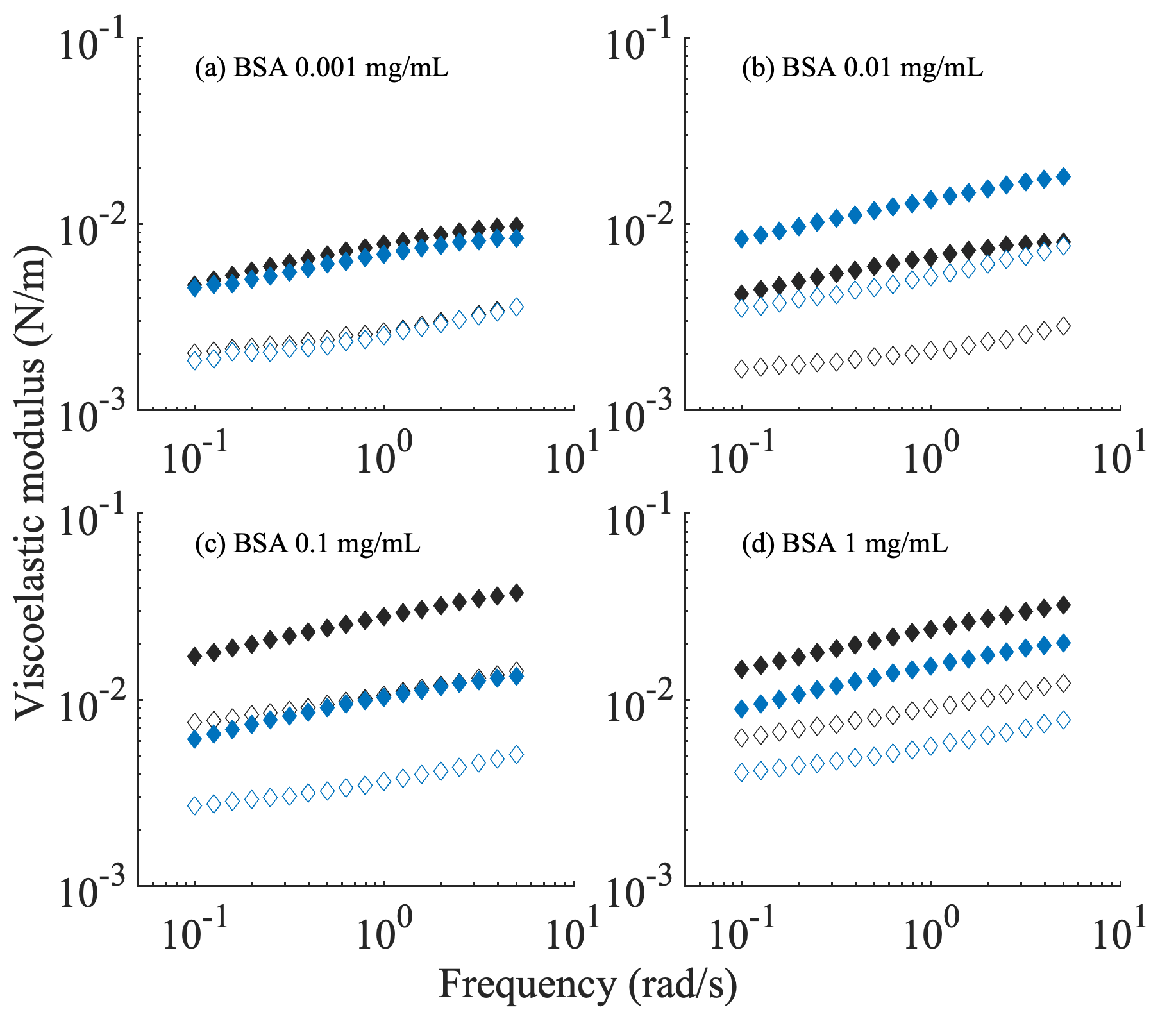

**Figure S-6**: Elastic and loss modulus as a function of angular frequency for BSA at concentrations of (a) 0.001 mg/mL, (b) 0.01 mg/mL, (c) 0.1 mg/mL, and (d) 1 mg/mL. For all concentrations, data without arginine (black markers) and with arginine (blue markers) are shown. Open markers represent the loss modulus, while solid markers correspond to the elastic modulus.
